# Design of biochemical pattern forming systems from minimal motifs

**DOI:** 10.1101/666362

**Authors:** Philipp Glock, Fridtjof Brauns, Jacob Halatek, Erwin Frey, Petra Schwille

**Affiliations:** Max-Planck-Institute of Biochemistry, D-82152 Martinsried; Arnold Sommerfeld Center for Theoretical Physics and Center for NanoScience, Department of Physics, Ludwig-Maximilians-Universität München, Theresienstraße 37, D-80333 München; Biological Computation Group, Microsoft Research, Cambridge CB1 2FB, UK

## Abstract

Although molecular self-organization and pattern formation are key features of life, only very few pattern-forming biochemical systems have been identified that can be reconstituted and studied *in vitro* under defined conditions. A systematic understanding of the underlying mechanisms is often hampered by multiple interactions, conformational flexibility and other complex features of the pattern forming proteins. Because of its compositional simplicity of only two proteins and a membrane, the MinDE system from *Escherichia coli* has in the past years been invaluable for deciphering the mechanisms of spatiotemporal self-organization in cells. Here we explored the potential of reducing the complexity of this system even further, by identifying key functional motifs in the effector MinE that could be used to design pattern formation from scratch. In a combined approach of experiment and quantitative modeling, we show that starting from a minimal MinE-MinD interaction motif, pattern formation can be obtained by adding either dimerization or membrane-binding motifs. Moreover, we show that the pathways underlying pattern formation are recruitment-driven cytosolic cycling of MinE and recombination of membrane-bound MinE, and that these differ in their *in vivo* phenomenology.

## Introduction

Patterns are a defining characteristic of living beings, and are found throughout all kingdoms of life. In the last years, it has become increasingly clear that protein patterns formed by reaction–diffusion mechanisms are responsible for a large range of spatiotemporal regulation (***Green and Sharpe, 2015***). Such processes allow organisms and cells to achieve robust intracellular patterning rooted in basic physical and chemical principles.

However, there is a lack of mechanistic understanding of the relationship between biomolecular features of proteins, i.e. their interaction domains and conformational states, and the collective properties of protein networks resulting in self-organized pattern formation. In other words, it is often unclear what exactly constitutes a *mechanism* of self-organization *on the biochemical level*. A major question is to what degree system-level biological functions, e.g. geometry sensing or length-scale selection, depend on particular biomolecular features. Some of these features may be essential for function, others may be irrelevant or redundant. The ability to unravel this *feature–function relationship* crucially depends on our ability to reconstitute biochemically distinct minimal systems experimentally and to compare these minimal variants to corresponding quantitative theoretical models. The key merit of such a combined approach is the ability to dissect different network architectures and also explore a broad range of reaction rates, and thereby uncover biomolecular mechanisms for system-level properties.

Here we address this feature-function relationship in the context of a fairly well-understood biological pattern-forming system: the Min-protein system of *Escherichia coli*. All its components are known – only two proteins are needed to form the pattern (MinD and MinE) – and the system has been successfully reconstituted in an easily malleable *in vitro* system (***Loose et al., 2008***; ***Ivanov and Mizuuchi, 2010***; ***Vecchiarelli et al., 2014***; ***Caspi and Dekker, 2016***; ***Kretschmer et al., 2017***). In the bacterial cell, this system contributes to the positioning of FtsZ, a key component of the division ring, at mid-cell. Two proteins, MinD and MinE, oscillate between the cell poles and thereby form a concentration gradient with a minimum at mid-cell. MinC, piggybacking on MinD, consequently inhibits FtsZ polymerization at the poles and thus positions the Z-ring in the middle.

Even though the Min protein system seems simple at first glance, there is much (and biologically relevant) complexity within the protein domain sequences and structures, and hence in the interaction between proteins. MinD is an ATPase which is believed to dimerize upon ATP-binding, raising its membrane affinity via the C-terminal membrane targeting sequence (MTS) (***Lackner et al., 2003***; ***Hu et al., 2002***; ***Szeto et al., 2003***). Bound to the membrane, MinD recruits further MinD-ATP, as well as its ATPase-activating protein MinE, which together form membrane-bound MinDE complexes (***Hu and Lutkenhaus, 2001***; ***Hu et al., 2002***). MinE stimulates MinD’s ATPase activity, thereby initiating disintegration of MinDE complexes and subsequent release of MinE and ADP-bound MinD into the cytosol. MinE, although only 88 amino acids in length, is a biochemically complex protein. It is found as a dimer in two distinct conformations (***Pichoff et al., 1995***; ***Park et al., 2011***): While diffusing in the cytoplasm, both the N-terminal MTS and the sequence directly interacting with MinD are buried within the protein. Upon sensing membrane-bound MinD, these features are released, which allows interaction with both the membrane and MinD (***Park et al., 2011***).

In summary, MinE exhibits four distinct functional features: activating MinD’s ATPase, membrane binding, dimerization, and a switch between an open, active and a closed, inactive conformation. The roles of these distinct functional features of MinE for pattern formation have previously been studied and discussed in the literature (***Vecchiarelli et al., 2016***; ***Kretschmer et al., 2017***; ***Denk et al., 2018***). It has been shown that MinE’s conformational switch is not essential for pattern formation, but conveys robustness to the Min system, as it allows pattern formation over a broad range of ratios between MinE and MinD concentrations (***Denk et al., 2018***). Furthermore, membrane binding of MinE was found to be non-essential for pattern formation (***Kretschmer et al., 2017***). These previous studies essentially retained the structure of MinE, predominantly mutating single residues.

Here, we chose a more radical strategy, in order to attempt a minimal design of fundamental modules towards protein pattern formation from the bottom-up. Specifically, we reduced MinE to its bare minimum function: binding to MinD, and thereby catalyzing MinD’s ATPase activity. We then reintroduced additional features—membrane binding and dimerization—one by one in a modular fashion, to study their specific role in pattern formation. This approach allowed us to identify the essential biochemical modules of MinE, and show that these facilitate two biochemically distinct mechanisms of pattern formation. We further analyzed these mechanisms in terms of reaction–diffusion models using theoretical analysis and numerical simulation. In particular, we show that the dimerization-driven mechanism is likely to be the dominant one for in *in vivo* pattern formation.

## Results and Discussion

Full flexibility and control over all parameters was achieved by reconstituting purified Min proteins and peptides in an *in vitro* well setup consisting of a glass-supported lipid bilayer with a large, open reservoir chamber (see Methods section for further details). To minimize the complexity of MinE in this reconstituted experimental system, we removed all sequences not in direct contact with MinD, keeping only 19 amino acids (13-31, further referred to as minimal MinE peptide) (***Figure 1***).

**Figure 1.**
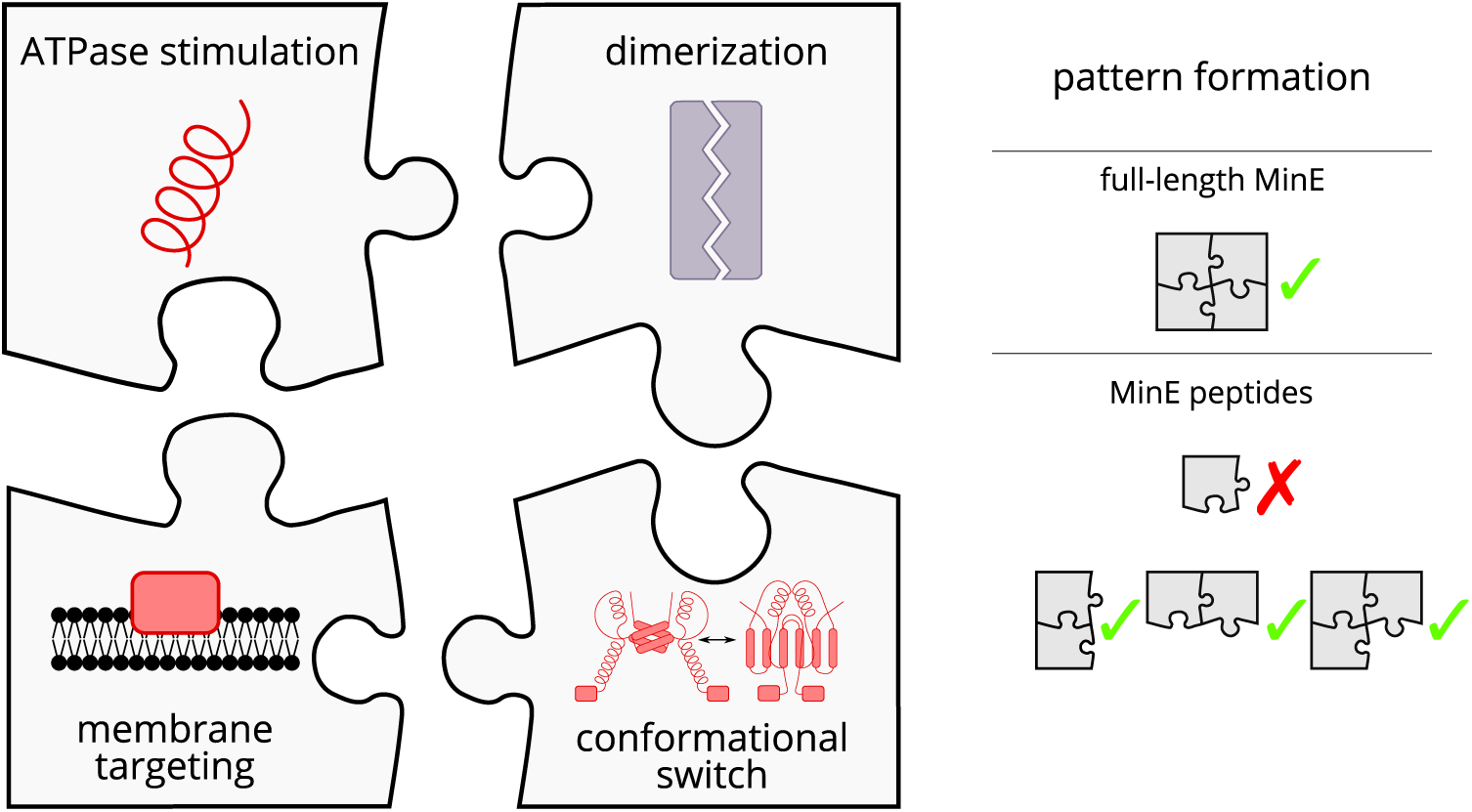
Schematic of the modular approach we took to engineering MinE in the *in vitro* Min system. While MinE has the core function to stimulate MinD’s ATPase, three additional properties help MinE to facilitate the emergence of spatiotemporal patterns. We show that two of these properties, dimerization and membrane targeting, can be modularly added to a minimal MinE peptide to facilitate pattern formation.

In agreement with previous studies, we observed (***Loose et al., 2008***; ***Glock et al., 2018b***) that the native *in vitro* Min system, consisting of MinD and full-length MinE, forms traveling (spiral) waves (see ***Figure 2***a) and (quasi-)stationary patterns. In contrast, we did not observe pattern formation for the reconstituted system containing the minimal MinE peptide in the nanomolar to low micromolar range (see ***Figure 2***b), suggesting that it lacks essential molecular features for pattern formation. Instead, membrane binding of MinD was dominant even for high concentrations of up to 20 µM of the minimal MinE peptide. We next tried to rescue pattern formation capability by re-introducing biomolecular features of MinE in a modular fashion.

**Figure 2.**
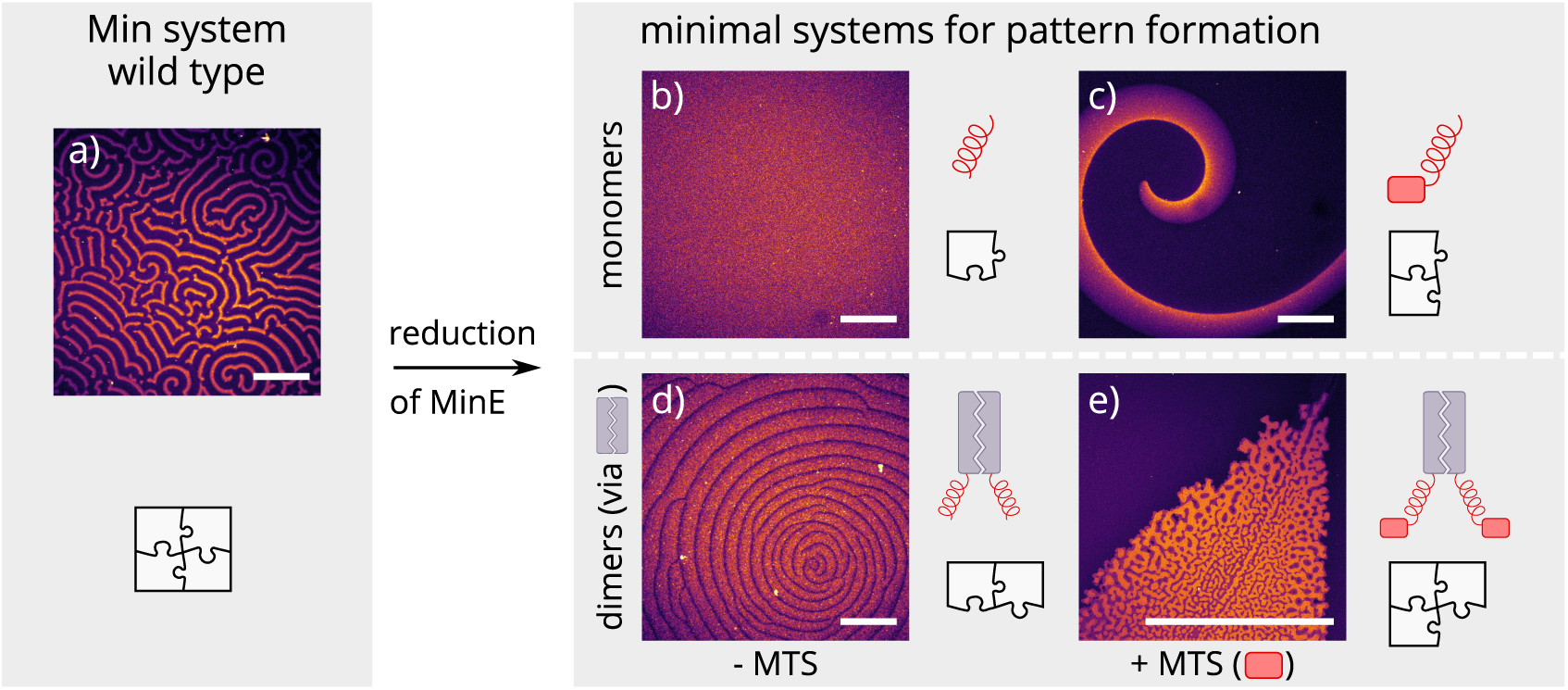
Patterns formed by the Min system and our minimal biochemical interaction networks. MinD and MinE self-organize to form evenly spaced travelling waves when reconstituted on flat lipid bilayers. Substituting MinE with minimal peptides leads to patterns if either a membrane-targeting sequence is added or the peptides are dimerized. Combining both approaches produces quasi-stationary patterns (top right), while a truly minimal peptide without either function cannot form patterns with MinD (d). (Concentrations and proteins used: (a) 1 µM MinD (always 70 % doped with 30 % Alexa647-KCK-MinD), 6 µM MinE-His; (b) 1.2 μM MinD, 50 nM MinE(13-31); (c) 1.2 µM MinD, 50 nM MinE(1-31); scalebars = 300 µm; (d) 1 µM MinD, 100 nM MinE(13-31)-Fos; (e) 1.2 μM MinD, 100 nM MinE(1-31)-GCN4.) **Figure 2–Figure supplement 1.** Global view of pattern formation by minimal systems **Figure 2–Figure supplement 2.** Titration results for MinE(1-31) and MinE(2-31)-sfGFP **Figure 2–video 1.** MinE(1-31) forms chaotic patterns with MinD. **Figure 2–video 2.** MinE(2-31)-msfGFP forms chaotic patterns with MinD.

Previous theoretical research has elucidated the key role of MinE cycling for the Min oscillations (***Halatek and Frey, 2012***). Each cycling step of MinE displaces one MinD from the membrane and thereby drives the oscillations that underlie pattern formation (***Halatek et al., 2018***). Specifically, in this model, MinE is assumed to cycle between a cytosolic state and a MinD-bound state on the membrane. To facilitate pattern formation, this cytosolic-cycling mechanism requires sufficiently strong recruitment of cytosolic MinE by membrane bound MinD (***Halatek and Frey, 2012***) suggesting that the recruitment rate of the minimal MinE peptide is too low. As the native MinE is a dimer, we hypothesized that dimerization might lead to increased recruitment, thus rescuing pattern formation. To test this hypothesis, we introduced dimerization back to the minimal MinE peptide by synthetically fusing it with well-described human and yeast leucine-zippers (Fos, Jun and GCN-4) (***Figure 1***) (***Szalóki et al., 2015***; ***O’Shea et al., 1989***). Indeed, this modification enabled sustained pattern formation in the system (see ***Figure 2***d). Compared to native MinDE patterns, those formed by dimerized peptides have larger wavelengths and are less coherent.

Another feature of native MinE that has been discussed in the context of pattern formation is persistent membrane binding via a membrane targeting sequence (MTS) (***Loose et al., 2011***). The MTS allows MinE to remain membrane-bound after its interaction with MinD, i.e. it decreases the detachment rate of MinE. MinE on the membrane can recombine with membrane-bound MinD. Hence, in addition to cycling between membrane-bound and cytosolic states, the MTS allows MinE to cycle between free and MinD-bound states on the membrane. This might alleviate the requirement for recruitment from the cytosol. To test whether the persistent membrane binding of MinE can facilitate pattern formation, we added back the MTS found in native MinE (residues 2-12) to the N-terminus of the peptide. This construct, contrary to published results (***Vecchiarelli et al., 2016***), forms patterns with MinD. As shown in ***Figure 2***c, the observed patterns are traveling waves with wavelengths several orders of magnitude larger than those found for the native *in vitro* Min system. Patterns are sustained over many hours within our assay.

Combining both features, i.e. adding both the MTS and a dimerization sequence to the minimal MinE peptide, resulted in (quasi-)stationary patterns, but the exact outcome depended heavily on the starting conditions of the assay (see ***Figure 2***e). In general, patterns formed by MinD and our minimal MinE peptides do not show the same degree of order as patterns formed by the wild-type Min proteins (***Glock et al., 2018b***) or MinD and His-MinE (***Loose et al., 2008***). In particular, there is no well-controlled characteristic length scale (wavelength), and the defined spirals or stationary patterns observed in the wild type Min system are sometimes replaced by chaotic centers as shown in ***Figure 2***d. The chaotic behavior is especially pronounced at high MinD concentrations (in this case with a minimal MinE plus MTS and sfGFP or MinE(1-31), respectively) (***video 1*** and ***video 2***).

Our experimental results suggest that two distinct features of MinE, dimerization and membrane binding, independently facilitate pattern formation of our reconstituted Min system with engineered, minimal MinE peptides. To support these conclusions and gain further insight into the mechanisms underlying pattern formation we performed a theoretical analysis using a reaction–diffusion model that captures all of the above biomolecular features. We extended the Min “skeleton” model introduced in (***Huang et al., 2003***; ***Halatek and Frey, 2012***) by MinE membrane binding, similar to the extension considered in (***Denk et al., 2018***). In this model, dimerization of MinE is effectively accounted for by an increased MinE recruitment rate. We performed linear stability analysis of the reaction–diffusion system to find the parameter regimes where patterns form spontaneously from a homogeneous initial state. The two-parameter phase diagram shown in ***Figure 3***(a) shows that increased MinE recruitment as well as slower MinE detachment can rescue pattern formation, in accordance with the *in vitro* experiments. This further supports the conclusion that the two biomolecular features of native MinE, namely the ability to dimerize and the MTS, facilitate pattern formation via two independent cycling pathways of MinE: cytosolic cycling and membrane recombination.

**Figure 3.**
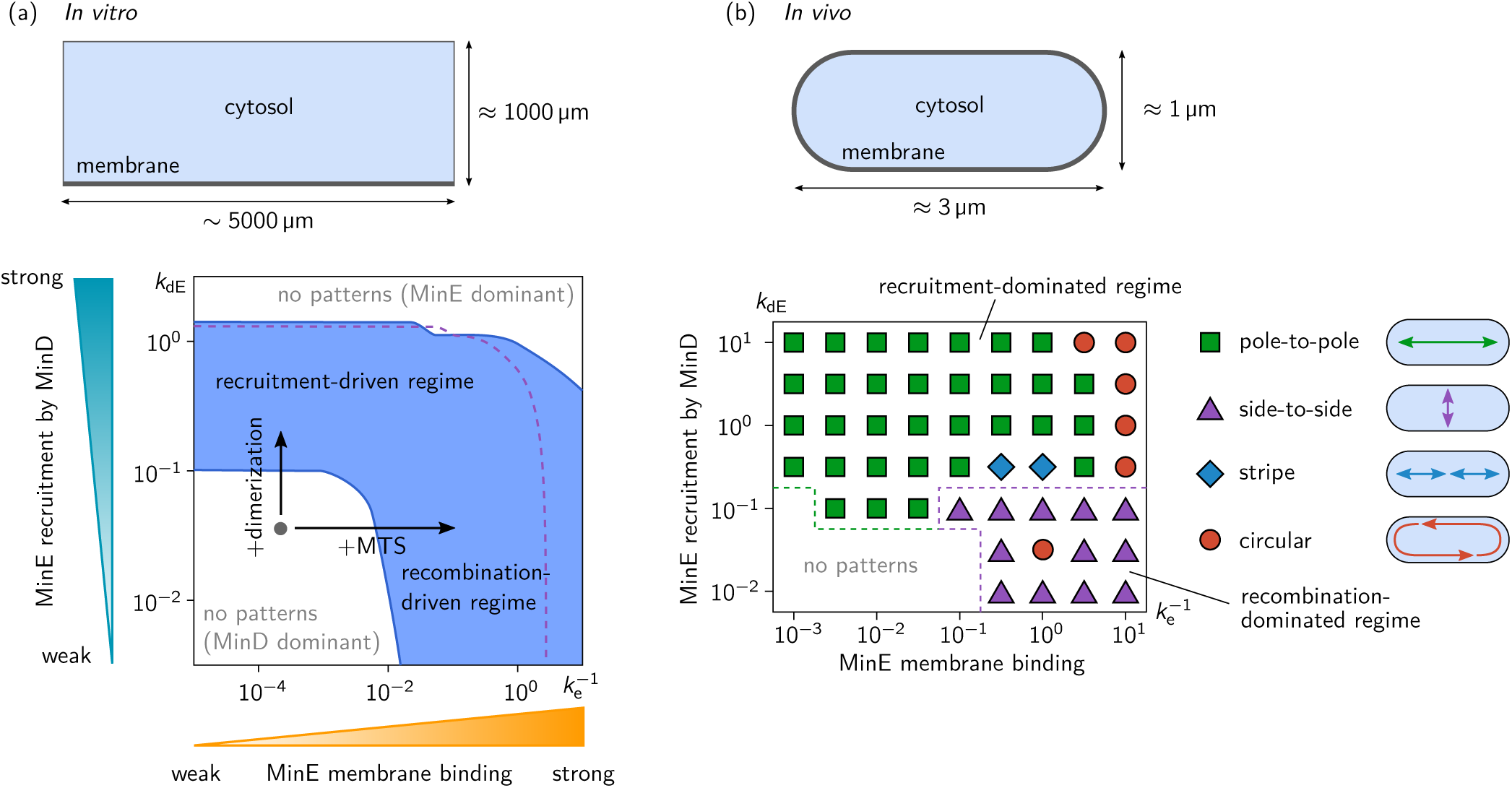
Pattern forming capability of the extended Min model (***Figure Supplement 1***) *in vitro* and *in vivo*. (a) *In vitro* geometry and two-parameter phase diagram obtained by linear stability analysis, showing the pattern formation capabilities of the MinDE-system in dependence of MinE membrane-binding strength 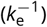 and MinE-recruitment rate *k*_dE_. The regime of spontaneous pattern formation (lateral instability) is indicated in blue The gray circle represents minimal MinE(13-31) construct, which does not facilitate self-organized pattern formation. The experimental domain additions are accounted for by respective changes of the kinetic rates, as indicated by the arrows. (Parameters: see ***section***; blue region: *k*_E_ = 0; purple dashed line: boundary of the pattern-formation regime for *k*_E_ = 5 µm^3^ s^*−*1^.) (b) Two-parameter phase diagram obtained by numerical simulations in *in vivo* geometry. We find regimes of different oscillation pattern types: pole-to-pole oscillations (green squares); side-to-side oscillations (purple triangles); stripe oscillations (blue diamonds); and circular waves (red circles). Videos 1-5 show examples each of these pattern types. **Figure 3–Figure supplement 1.** Network cartoon of the MinE “skeleton” model extended by MinE membrane binding. **Figure 3–Figure supplement 2.** Linear stability analysis in the ellipse geometry. **Figure 3–video 1.** Pole-to-pole oscillation for weak MinE binding (*k*_dE_ = 3.16 µm^3^s^−1^, *k*_e_ = 1000 s^−1^). **Figure 3–video 2.** Pole-to-pole oscillation for strong MinE binding (*k*_dE_ = 3.16 µm^3^s^−1^, *k*_e_ = 0.316 s^−1^). **Figure 3–video 3.** Circular wave (*k*_dE_ = 3.16 µm^3^s^−1^, *k*_e_ = 0.1 s^−1^). **Figure 3–video 4.** Stripe oscillation (*k*_dE_ = 0.316 µm^3^s^−1^, *k*_e_ = 3.16 s^−1^). **Figure 3–video 5.** Side-to-side oscillation (*k*_dE_ = 0.1 µm^3^s^−1^, *k*_e_ = 0.316 s^−1^).

To test whether either or both of these two pattern-forming pathways fulfill the biological function of the Min-protein patterns, we studied pattern formation using the generalized reaction– diffusion model taking into account realistic cell geometry. In *E. coli*, Min oscillations have to take place along the long axis of the rod-shaped cells for correct positioning FtsZ at midcell. Interestingly, linear stability analysis (see ***Figure 3–Figure Supplement 2***) shows that the membrane-recombination driven mechanism favors short-axis oscillations which is at odds with the biological function of the Min system. Indeed, our numerical simulations show that pole-to-pole oscillations are only possible for sufficiently strong cytosolic cycling, whereas the recombination-driven mechanism leads to side-to-side oscillations (see ***Figure 3***(b)). A recent theoretical study on axis-selection of the PAR system in *C. elegans* suggests that pattern formation driven by an antagonism of membrane bound proteins generically leads to short-axis selection (***Gessele et al., 2018***). Here, membrane-bound MinE antagonizes membrane-bound MinD via the membrane-recombination pathway. sufficiently strong MinE-recruitment from the cytosol supersedes the membrane-recombination pathway and leads to long-axis selection (pole-to-pole oscillations) even when MinE-membrane binding is strong.

Taken together, we conclude that Min-pattern formation *in vivo* is driven by cytosolic cycling of MinE, because correct axis selection (pole-to-pole oscillations) is essential for cell-division of *E. coli* and other gram-negative bacteria. In a broader context, our results demonstrate that multiple mechanisms with different characteristics, e.g. in their ability to sense geometry, can coexist in one reaction network. Most importantly, this highlights that a classification of pattern-forming mechanisms in terms of the reaction network topology alone misses important aspects of pattern formation that can be crucial for the biological function.

With respect to a potential biochemical origin of the pattern-forming mechanisms, we showed how additional protein domains can move the whole system into a mechanistically distinct regime. Enhancing the strength of MinE recruitment by MinD via dimerization shifts the system into a regime of recruitment-driven pattern formation. Alternatively, adding membrane targeting to the peptide unlocked a new pathway and led to sustained patterns via MinD-MinE recombination on the membrane (see supplementary discussion in ***Appendix 1*** for further details).

In conclusion, the concept of modular engineering of pattern formation through distinct protein domains adds an entirely new dimension to the Min system, and establishes it further as a paradigmatic model for studying the mechanisms underlying self-organized pattern formation. Now, defined modules can be added, removed and interchanged. Interestingly, our experimental findings provide evidence that the distinct functional modules of MinE need not be provided by native parts of the proteins, but can be substituted with foreign sequences. Moreover, the part of MinE that interacts with MinD can be added as a small peptide tag of 19 amino acids to any host protein (as shown for superfolder-GFP + MTS, ***Figure Supplement 2***), leading to a chimera protein that inherits key properties, such as membrane-interactions and protein-protein interactions, from the host protein. The modular domains provide an experimental platform to systematically modify the molecular interactions. Together with systematic theoretical studies, this is a powerful and versatile tool to study the general principles underlying biological pattern formation in multispecies, multicomponent reaction–diffusion systems.

## Methods and Materials

Most experimental methods used in this publication were exhaustively described in text and video in a recent publication (***Ramm et al., 2018***). We therefore describe these techniques only in brief.

### Membranes

SLBs were prepared from DOPC and DOPG (ratio 2:1) small unilamellar vesicles in Min buffer (25 mM Tris-HCl pH 7.5, 150 mM KCl, 5 mM MgCl_2_) by adding them (at 0.53 mg/mL) on top of a charged, cleaned glass surface. The solution was diluted after one minute by addition of 150 mL Min buffer. After a total of 3 minutes, membranes in chambers were washed with 2 mL of Min buffer.

### Assay chamber

Assay chambers were assembled from piranha-cleaned coverslips and a cut 0.5 ml plastic reaction tube by gluing the tube upside down onto the cleaned and dried surface using UV-curable adhesive.

### *In vitro* self-organization assay

The buffer volume in an assay chamber containing an SLB was adjusted to yield a final volume of 200 µL including protein solutions and ATP. Proteins, peptides and further reactants were added and the solution was mixed by pipetting.

### Peptides

Peptides were synthesized using Fmoc chemistry by our in-house Biochemisty Core Facility. MinE(2-31)-KCK-Atto488 was expressed as a SUMO fusion in *E. coli* BL-21 DE3 pLysS cells, the SUMO tag was then cleaved using SenP2 protease and the remaining peptide was labelled using Atto488-maleimide to site-Specifically target the cysteine residue. Labelling was done as described below.

### Protein design and purifications

Detailed information about cloning procedures and design of proteins can be found in the supplementary information.

### Protein concentration measurements

Protein concentrations were determined by using a modified, linearized version of the Bradford assay in 96-well format (***Ernst and Zor, 2010***).

### Labelling

Atto 488-maleimide in 5-7 µL DMSO (about three molecules of dye per protein) was added dropwise to *∼* 0.5 mL of protein solution in storage buffer (50 mM HEPES pH 7.25, 300 mM KCl, 10 % glycerol, 0.1 mM EDTA, 0.4 mM TCEP) in a 1.5 mL reaction tube. The tube was wrapped in aluminium foil and incubated at 4 °C on a rotating shaker for two to three hours. Free dye was separated from proteins first by running the solution on a PD-10 buffer exchange column equilibrated with storage buffer. Then, remaining dye was diluted out by dialysis against storage buffer overnight. The labelling efficiency was measured by recording an excitation spectrum of the labelled protein and measuring the protein concentration as described above. We then calculated the resulting labelling efficiency using the molar absorption provided by the dye supplier (Atto 488: 9.0 *×* 10^4^ M^*−*1^ cm^*−*1^).

### Imaging

Microscopy was done on commercial Zeiss LSM 780 microscopes with 10x air objectives (Plan-Apochromat 10x/0.45 M27 and EC Plan-Neofluar 10x/0.30 M27). Tile scans with 25 tiles (5×5) at zoom level 0.6 were stitched to obtain overview images of entire assay chambers and resolve the large-scale patterns formed. More detailed images and videos were acquired on the same instruments using EC Plan-Neofluar 20x/0.50 M27 or Plan-Apochromat 40x/1.20 water-immersion objectives.

### The Min “skeleton model” extended by MinE membrane binding

To capture the effect of MinE membrane binding, we extend the “skeleton” model introduced in (***Halatek and Frey, 2012***). ***Figure 3–Figure Supplement 1*** shows a cartoon of the reaction network. We present the model first for a general geometry with a cytosolic volume coupled to a membrane surface. To perform linear stability analysis, we implemented this model in a “box geometry” representing the *in vitro* setup with a membrane at the bottom, and in an ellipse geometry mimicking the rod-like cell shape of *E. coli*.

On the membrane, proteins diffuse and undergo chemical reactions, including attachment, detachment and interactions between membrane-bound proteins

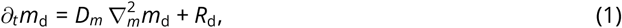

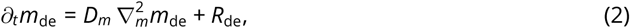

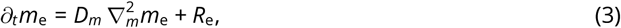

where ∇_*m*_ is the gradient operator along the membrane. In the cytosol, proteins diffuse and MinD undergoes nucleotide exchange with a rate *λ*

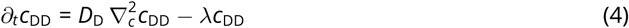

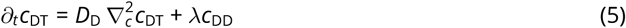

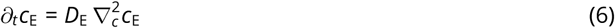

The two domains are coupled via the boundary conditions at the membrane

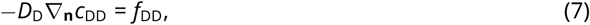

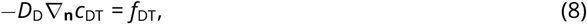

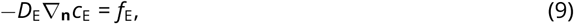

where ∇**_n_** is the gradient along the inward pointing normal to the membrane. The reaction terms are derived from the interaction network ***Figure 3–Figure Supplement 1*** via the mass-action law and read

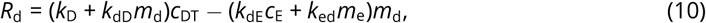

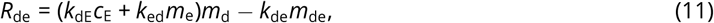

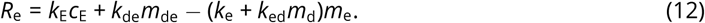

Correspondingly, the attachment-detachment flows are

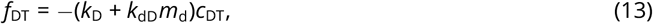

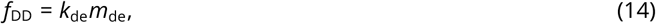

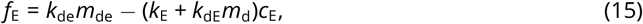

such that the dynamics conserve the global total densities of MinD and MinE

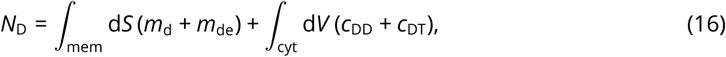

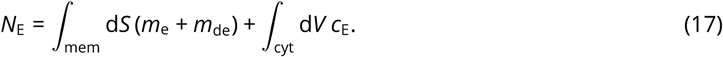

### Linear stability analysis

To perform linear stability analysis, we need to find a set of orthogonal basis functions that fulfill the boundary conditions and diagonalize the Laplace operator, ∇^2^, on both domains (membrane and cytosol) simultaneously. In general, this is not analytically possible in arbitrary geometry. However, in a box geometry with a flat membrane, a closed form of the basis functions can easily be obtained. Furthermore, in a two dimensional ellipse geometry, a perturbative ansatz can be used to obtain an approximate set of basis functions, as was shown in (***Halatek and Frey, 2012***) and used in (***Wu et al., 2016***; ***Gessele et al., 2018***). In the following, we briefly outline how the basis functions can be determined and employed to perform linear stability analysis. For details we refer to the supplementary materials of Refs. (***Halatek and Frey, 2018***; ***Denk et al., 2018***; ***Halatek and Frey, 2012***; ***Gessele et al., 2018***).

#### *In vitro* box geometry

For linear stability analysis of the *in vitro* system, we consider a two-dimensional box with a membrane at the bottom surface, representing a slice through the *in vitro* system. The cytosol domain is a rectangle in the *x*-*z* plane with height *h* and length *L*. The bottom boundary at *z* = 0 is the one-dimensional membrane domain – a line of length *L*. It is coupled to the bulk via reactive boundary conditions, Eqs. (7) to (9). The other boundaries of the rectangular bulk domain are equipped with reflective boundaries. In this geometry, the gradient operators tangential and normal to the membrane are simply ∇_*m*_ ≡ *∂*_*x*_ and ∇**_n_** *≡ ∂_z_*.

The first step of a linear stability analysis is to calculate the steady state whose stability is to be analyzed. Typically this is a homogeneous steady state. In the system considered here, the most simple steady state is homogeneous along the *x*-direction. However, there must be cytosolic gradients in the *z*-direction due to the reactive boundary condition and the nucleotide exchange in the cytosol. Because the cytosol dynamics are linear, they can be solved in closed form.

To analyze the stability of such a steady state, one linearizes the dynamics around it. The ansatz to solve the resulting linear system is to diagonalize the Laplace operator. Importantly, in a system with multiple coupled domains, one needs to find a set of basis functions that diagonalize the Laplace operator on all domains (here membrane and cytosol), and that fulfill the reactive boundary conditions that couple these domains, simultaneously. In the *x*-direction, i.e. the lateral direction along the one-dimensional membrane, the eigenfunctions are simply Fourier modes. The bulk eigenfunctions in the *z*-direction, normal to the membrane, are exponential profiles and can be obtained in closed form by solving the linear cytosol dynamics, Eqs. (4) to (6).

These eigenfunctions can then be plugged into the the membrane dynamics and the boundary conditions linearized around the homogeneous steady state. The resulting set of linear algebraic equations can be solved for the growth rates of the Fourier modes. Thus, one obtains a relationship between wavenumber *q* of a mode and its growth rate *σ*(*q*). This relationship is called dispersion relation.

For details of the implementation of the linear stability analysis outlined above, we refer the reader to the supplementary materials of (***Halatek and Frey, 2018***) and (***Denk et al., 2018***).

#### *In vivo* ellipse geometry

Linear stability analysis in an ellipse geometry is technically more involved, because the curved boundary makes it impossible to find a common eigenbasis of the Laplace operator on membrane and cytosol in closed form. For a detailed exposition of linear stability analysis in an elliptical geometry, we refer the reader to the supplementary materials of (***Halatek and Frey, 2012***).

### Parameters

#### In vitro

We used the same parameters as in (***Halatek et al., 2018***): *k*_D_ = 0.065 µm s^*−1*^, *k*_dD_ = 0.098 µm^2^ s^*−1*^, *k*_dE_ = 0.126 µm^2^ s^*−1*^, *k*_de_ = 0.34 µm s^*−1*^, *D*_*m*_ = 0.013 µm^2^ s^*−1*^, *D*_*D*_ = 16 µm^2^ s^*−1*^, *D*_*E*_ = 10 µm^2^ s^*−1*^, *n*_E_ = 120 µm^*−*3^, *n*_D_ = 1200 µm^*−*3^.

Additionally, we set the MinE membrane recombination rate to *k*_ed_ = 0.1 µm s^*−*1^, and varied the MinE detachment rate, *k*_e_, in the range 10^*−*1^ µm s^*−*1^ to 10^5^ µm s^*−*1^. To test the effect of spontaneous MinE membrane attachment (*k*_E_ *>* 0) we compared the results from LSA for *k*_E_ = 0 and *k*_E_ = 10 µm^3^ s^*−*1^, and found that spontaneous attachment is only relevant for very small MinE detachment rate, *k*_e_, i.e. strong MinE membrane binding, where it suppresses pattern formation due to a dominance of membrane-bound MinE (see ***Figure 3***(a), purple dashed line).

Note that the bulk height dependence saturates above around 50 µm, the maximal penetration depth of bulk gradients (***Halatek and Frey, 2018***). The bulk heights in the experiments were well above this saturation threshold at around 1 mm, allowing us to use the limit of large bulk height *h*.

#### In vivos

We used the same parameters as in (***Halatek and Frey, 2012***): *k*_D_ = 0.1 µm s^*−1*^, *k*_dD_ = 0.108 µm^2^ s^*−1*^, *k*_dE_ = 0.65 µm^2^ s^*−1*^, *k*_de_ = 0.4 µm s^*−1*^, *D*_*m*_ = 0.013 µm^2^ s^*−1*^, *D*_*D*_ = 16 µm^2^ s^*−*1^, *D*_*E*_ = 10 µm^2^ s^*−*1^, *N*_E_ = 700, *N*_D_ = 2000.

Additionally, we set the MinE membrane recombination rate to *k*_ed_ = 0.2 µm s^*−*1^, and varied the MinE detachment rate, *k*_e_, in the range 10^−1^ µm s^*−*1^ to 10^3^ µm s^*−*1^. As in the *in vitro* case, spontaneous MinE membrane attachment (*k*_E_ *>* 0) has no significant effect, so we set *k*_E_ = 0. (LSA and numerical simulations for *k*_E_ = 10 µm^3^ s^*−*1^ yield a phase diagram with the same qualitative structure as the one presented in ***Figure 3***(b).)

We mimic the cell geometry by an ellipse with lengths 0.5 µm and 2 µm for the short and long half axis respectively.

### Numerical simulations

The bulk-boundary coupled reaction–diffusion dynamics Eqs. (1) to (15) were solved using a finite element solver code (COMSOL Multiphysics).

Due to its large size, simulations of the *in vitro* system are very time consuming and beyond the scope of this work. Because most of the kinetic rates are not known, extensive parameter studies would be necessary to gain insight from such simulations.

## Acknowledgments

We thank the MPIB Biochemistry Core Facility, especially Stefan Pettera, for synthesizing peptides used in this study. We also thank Simon Kretschmer for stimulating discussions. PG acknowledges support by the International Max-Planck Research School for Molecular Life Sciences (IMPRS-LS). P.G. and F.B. acknowledge financial support by the DFG Research Training Group GRK2062 (‘Molecular Principles of Synthetic Biology’).

## Appendix 1

### Protein design and cloning

Several instances of MinE(2-31)-sfGFP were cloned, expressed and tested. We started with a construct carrying a His-tag on the N-terminus His-(MinE-2-31)-sfGFP. Then, we became concerned about dimerization of the fluorescent protein and introduced a mutation (V206K) to make His-MinE(2-31)-msfGFP. Then, we discovered that N-terminal tagging influences the properties of our minimal constructs and wt MinE and changed the construct to carrying a C-terminal His-tag (MinE(1-31)-msfGFP-His). The methionine residue was re-introduced here as a start codon, and is cleaved in E. coli. Additionally, we prepared MinE(13-31)-sfGFP and confirmed that without MTS, no patterns are formed.

The first construct, His-MinE(2-31)-sfGFP was cloned as follows: A fragment containing the pET28a vector-backbone and the start of His-MinE was amplified from pET28a-His-MinE using primers PG073+PG074. The sfGFP fragment was amplified from pVRB18-XX-sfGFP using primers PG069+PG070. The two fragments were recombined in E. coli to yield pET28a-His-MinE(2-31)-sfGFP. His-MinE(13-31)-sfGFP was assembled from three fragments. The sfGFP fragment was generated as described above. A second fragment containing the vector backbone and compatible overhangs was generated from pET28a-His-MinE using primers PG073+PG077. Finally, the MinE(13-31) fragment was amplified from pET28a-His-MinE using primers PG072+PG016, then a second PCR reaction was run on this fragment with primers PG076+PG074. All three fragments were recombined in E. coli.

His-MinE(2-31)-msfGFP was generated from His-MinE(2-31)-sfGFP by recombining two fragments generated by PCR with primers PG087+PG043 and PG088+PG044, respectively.

MinE(1-31)-msfGFP-His was recombined from two fragments. the MinE(1-31)-msfGFP was amplified from pET28a-His-MinE(2-31)-msfGFP using primers PG090+PG091. The vector fragment was generated from pET28a-BsMTS-mCherry-His using primers PG089+PG007.

Custom DNA sequences were ordered for GCN-4, c-Jun and c-Fos. DNA fragments consisting of a linker sequence, the respective leucine zipper and another linker sequence were amplified via PCR using primers PG103+PG104 (GCN4), PG105+PG106 (Jun) or PG107+PG108 (Fos). Similarly, FKBP and FRB were amplified using primers PG110+PG111 (FKBP) and PG112+PG113 (FRB). A fragment of MinE(13-31) containing compatible overlaps was generated from PCR on pET28a-MinEL-msfGFP-His using primers PG109+PG102. The vector containing MinE(1-31) and compatible overhangs was amplified from pET28a-MinE-His using primers PG007 and PG102. For the three-fragment assemblies, the vector was created via PCR from BsMTS-mCherry-His (Ramm et al.) using primers PG007+PG089. The desired construct vectors were then created via three-fragment homologous recombination in E. coli TOP10, or two-fragment in case of MinE(1-31) constructs. In an additional step, the protein sequence KCK was inserted into the MinE(13-31) constructs by amplifying two halves of the vector. The first half was amplified using primers PG114+PG43, the second half using primers PG115+PG44. After DpnI digest (done for all fragments amplified from functional vectors), the fragments were transformed in to E. coli TOP10 and selected on kanamycin LB plates for homologous recombination. All constructs’ integrity was verified via Sanger sequencing.

SUMO-MinE(1-31)-KCK-His and SUMO-MinE(13-31)-KCK-His were generated via homologous recombination of two fragments each. For the construct with MTS, one fragment was amplified from pET28M-SUMO1-GFP using primers PG043+PG116. The second fragment was amplified from pET28M-SUMO1-MinE (***Glock et al., 2018b***) using primers PG044+PG117. Fragments for the construct without MTS were amplified from the recombined vector described above using primers PG043+PG118 and from pET28M-SUMO1-GFP using primers PG044+PG119.

### Purification of proteins

MinD, MinD-KCK-Alexa647, mRuby3-MinD, His-MinE and MinE-His were purified as previously described (***Ramm et al., 2018***; ***Glock et al., 2018a***,b). MinE(13-31)-Fos, MinE(13-31)-Jun and MinE(13-31)-GCN4 were purified as described for MinE-His (***Glock et al., 2018b***). MinE(2-31)-Fos, MinE(2-31)-Jun and MinE(2-31)-GCN4 were highly insoluble and therefore entirely found in the pellet fraction after cell lysis and centrifugation. The supernatant was discarded and the pellet re-solubilised in lysis buffer U (8M Urea, 500 mM NaCl, 50 mM Tris-HCl pH 8) by pipetting, vortexing and submerging the vial in a sonicator bath. The residual insoluble fraction was pelleted by centrifugation at 50000 g for 40 minutes. The supernatant was incubated with Ni-NTA agarose beads (*∼* 2 mL per 400 mL initial culture) for 1 h at room temperature on a rotating shaker. Agarose beads were pelleted at 400 g, 4 min and the supernatant was discarded. Purification was continued at RT since proteins were unfolded and kept in 8 M Urea. Agarose beads were loaded on a glass column and washed three times with 10 mL of above lysis buffer U. Further washes (3x) were performed with wash buffer U (8 M Urea, 500 mM NaCl, 20 mM imidazole, 50 mM Tris-HCl pH 8). The protein was eluted with elution buffer U (8 M Urea, 500 mM NaCl, 300 mM imidazole, 50 mM Tris-HCl pH8) and fractions with the highest protein content (Bradford, by eye) were pooled. Re-folding of the pooled eluate was done by dialyzing in multiple steps. In a first step, the solution was dialyzed against buffer D1 (6 M Urea, 500 mM NaCl, 50 mM Tris-HCl pH 8, 10% glycerol) over night. In a second step, against buffer D2 (4 M Urea, 500 mM NaCl, 50 mM Tris-HCl pH8, 10% glycerol) for 2 h, then against buffer D3 (2 M Urea, 500 mM NaCl, 50 mM Tris-HCl pH8, 10% glycerol) for further 2 h. The final dialysis was done against storage buffer (300 mM KCl, 50 mM HEPES pH 7.25, 10% glycerol, 1 mM TCEP, 0.1 mM EDTA). To separate the re-folded protein from aggregates, the protein solution was ultracentrifuged for 40 min at 50000 g, 4 °C. Protein concentration was then determined as described in the methods section. MinE(13-31)-KCK-His-Atto 488 and MinE(2-31)-KCK-His-Atto 488 were expressed and purified as described for MinE-His. SUMO-peptide fusions were then added into 1:100 (protease:protein) of SenP2 protease and dialyzed against storage buffer. Labelling was performed as described in the methods section.

### Supplementary discussion

Going forward, it will be interesting to explore the Min system further along the avenue of individual protein domains / features and their role for self-organized pattern formation. We suspect that the minimization of MinE peptides could be taken even further by shortening the peptide. Especially at the C-terminus we expect that several residues do not contribute to function, since they are not visible in a crystal structure of MinE(13-31) with MinD (***Park et al., 2011***). Additionally, the peptide still retains residues required for the dual function in the context of the MinE switch. Therefore, an optimized and further reduced peptide could be screened for. Additionally, our experiments with minimal peptides added to a superfolder–GFP (***Figure Supplement 2***) show that unrelated proteins can be attached. This opens the possibility to couple the spatiotemporal pattern to a different protein system. In principle, any protein can act as a minimal MinE if a peptide can be added internally or at either terminus of the protein.

Although we have not tested this prediction, we expect that the native MTS of MinE could be replaced with another MTS in our minimal peptides to restore pattern formation. It would be interesting to exchange the native MTS for a quantitatively described, diverse set of MTS to determine the required strength of membrane anchors needed for minimal MinE pattern formation. However, no such set or even just quantitative data on binding strength of multiple MTS is available at the moment.

Since we relate the lack of pattern formation to the recruitment rate of MinE(13-31), it may be possible to alter MinE recruitment by changing the buffer conditions such as salt concentration, type of ions (e.g. Sodium instead of Potassium), viscosity or pH. We can only speculate here, however, since screening a vast amount of conditions was not in the scope of the present study. Studies done on the wild type Min system using different conditions showed some impact on pattern formation (***Downing et al., 2009***; ***Vecchiarelli et al., 2014***).

### Primers used in this study

PG007: AC-pET_for GTCGAGCACCACCACCA

PG016: B-pET-MinE_rev GTGCGGCCGCAAGCTTTTAGCGACGGCGTTCAGCAA

PG043: mut_KanR_fw TGAAACATGGCAAAGGTAGCGT

PG044: mut_KanR_rev GCTACCTTTGCCATGTTTCAGAAA

PG073: sfGFP-pET_fw CATGGATGAGCTCTACAAATAAAAGCTTGCGGCCGCAC

PG074: sfGFP-li-MinE31_rev AAAGTTCTTCTCCTTTGCTCACAGAACCAGAAGAACCAGAAGAGCGACGGCGTTCAGCAAC

PG075: sfGFP-MinE_L_fw CATGGATGAGCTCTACAAAGCATTACTCGATTTCTTTCTCTCGC

PG076: E-pET-MinEs_fw GGGTCGCGGATCCGAATTCAAAAACACAGCCAACATTGCAA

PG077: lolipET_rv GAATTCGGATCCGCGACC

PG087: sfGFP_V206K_fw TACCTGTCGACACAATCTAAGCTTTCGAAAGATCCCAAC

PG088: sfGFP_V206K_rev GTTGGGATCTTTCGAAAGCTTAGATTGTGTCGACAGGTA

PG089: pET28a-start_rev CATGGTATATCTCCTTCTTAAAGTTAAACAA

PG090: pET-MinEL_fw TAAGAAGGAGATATACCATGGCATTACTCGATTTCTTTCTCTCGC

PG091: pET-msfGFP_rev TGGTGGTGGTGGTGCTCGACTCCAGATCCACCTTTGTAGAGCT

PG103: GCN4_fw TCTTCTGGTTCTTCTGGTTCTCGTATGAAACAGCTGGAAGACAA

PG104: GCN4_rev GTGCTCGACTCCAGATCCACCACGTTCACCAACCAGTTTTTTC

PG105: Jun_fw TCTTCTGGTTCTTCTGGTTCTCGTATCGCTCGTCTGGAAGA

PG106: Jun_rev GTGCTCGACTCCAGATCCACCGTAGTTCATAACTTTCTGTTTCAGCTG

PG107: Fos_fw TCTTCTGGTTCTTCTGGTTCTCTGACCGACACCCTGCAG

PG108: Fos_rev GTGCTCGACTCCAGATCCACCGTAAGCAGCCAGGATGAATTCC

PG109: pET-MinEs_fw TAAGAAGGAGATATACCATGAAAAACACAGCCAACATTGCAAAAG

PG110: FKBP_fw TCTTCTGGTTCTTCTGGTTCTGGTGTTCAGGTCGAAACTATCTCTC

PG111: FKBP_rev GTGCTCGACTCCAGATCCACCTTCCAGTTTCAGCAGTTCAACG

PG112: FRB_fw TCTTCTGGTTCTTCTGGTTCTGAAATGTGGCATGAGGGTCTC

PG113: FRB_rev GTGCTCGACTCCAGATCCACCCTGTTTAGAGATGCGACGAAAGAC

PG114: li-KCK_fw GGATCTGGAGTCGAGAAATGCAAACACCACCACCACCAC

PG115: li-KCK_rev GTGGTGGTGGTGGTGTTTGCATTTCTCGACTCCAGATCC

PG116: KCK-pET_fw GTTCTTCTGGTAAATGCAAATGAAAGCTTGCGGCCG

PG117: EL_li_rev TTTGCATTTACCAGAAGAACCAGAACCGCGACGGCGTTCAGC

PG118: SUMO-Es-fw ACCAGGAACAAACCGGTGGATCAAAAAACACAGCCAACATTGCAAA

PG119: SUMO_rev TCCACCGGTTTGTTCCTGG

---

**Figure 2–Figure supplement 1.**
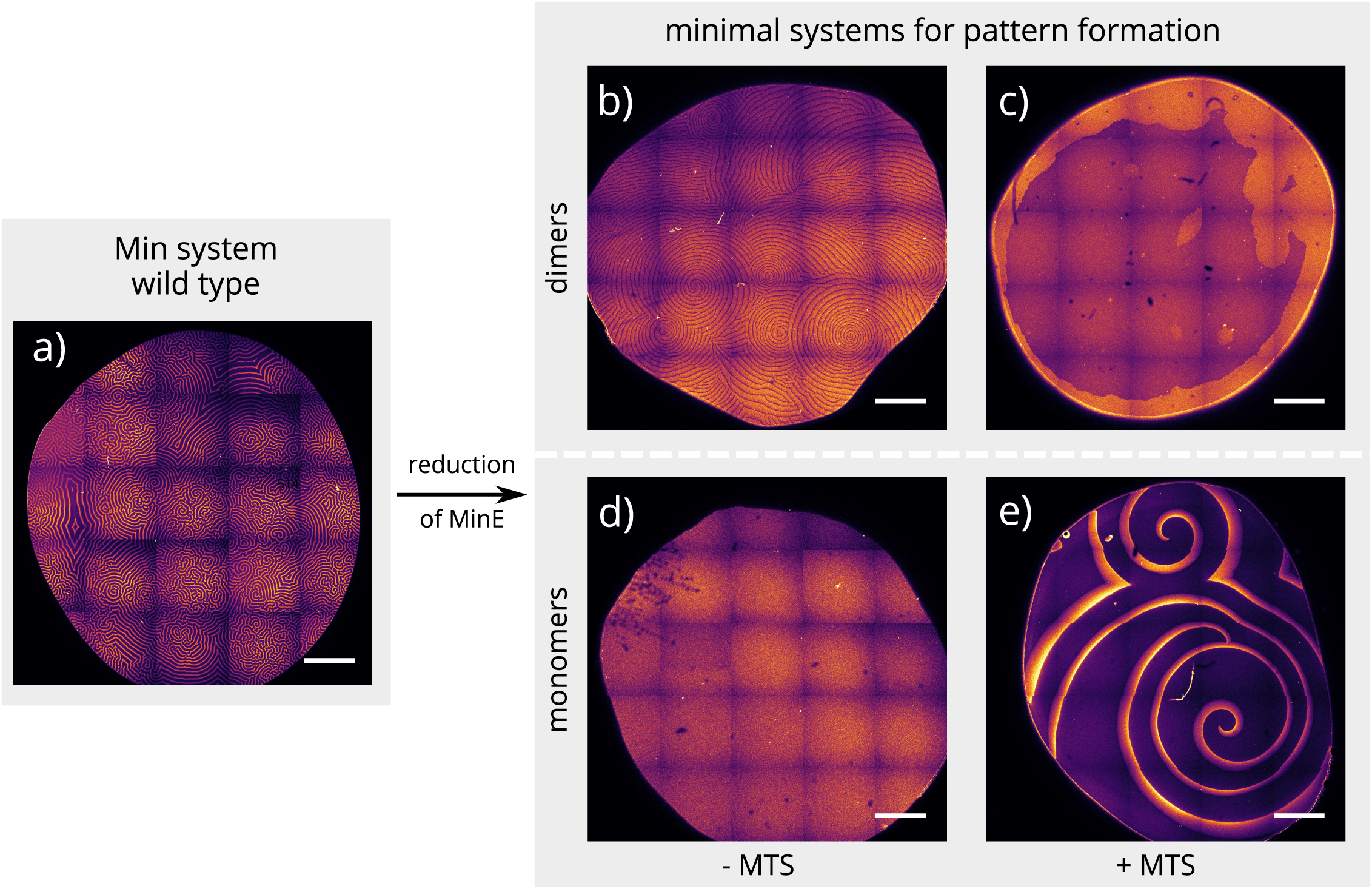
Overview images of the same experiment chambers as in Figure 2. (Concentrations and proteins used same as in main figure; scalebars = 1000 µm.)

**Figure 2–Figure supplement 2.**
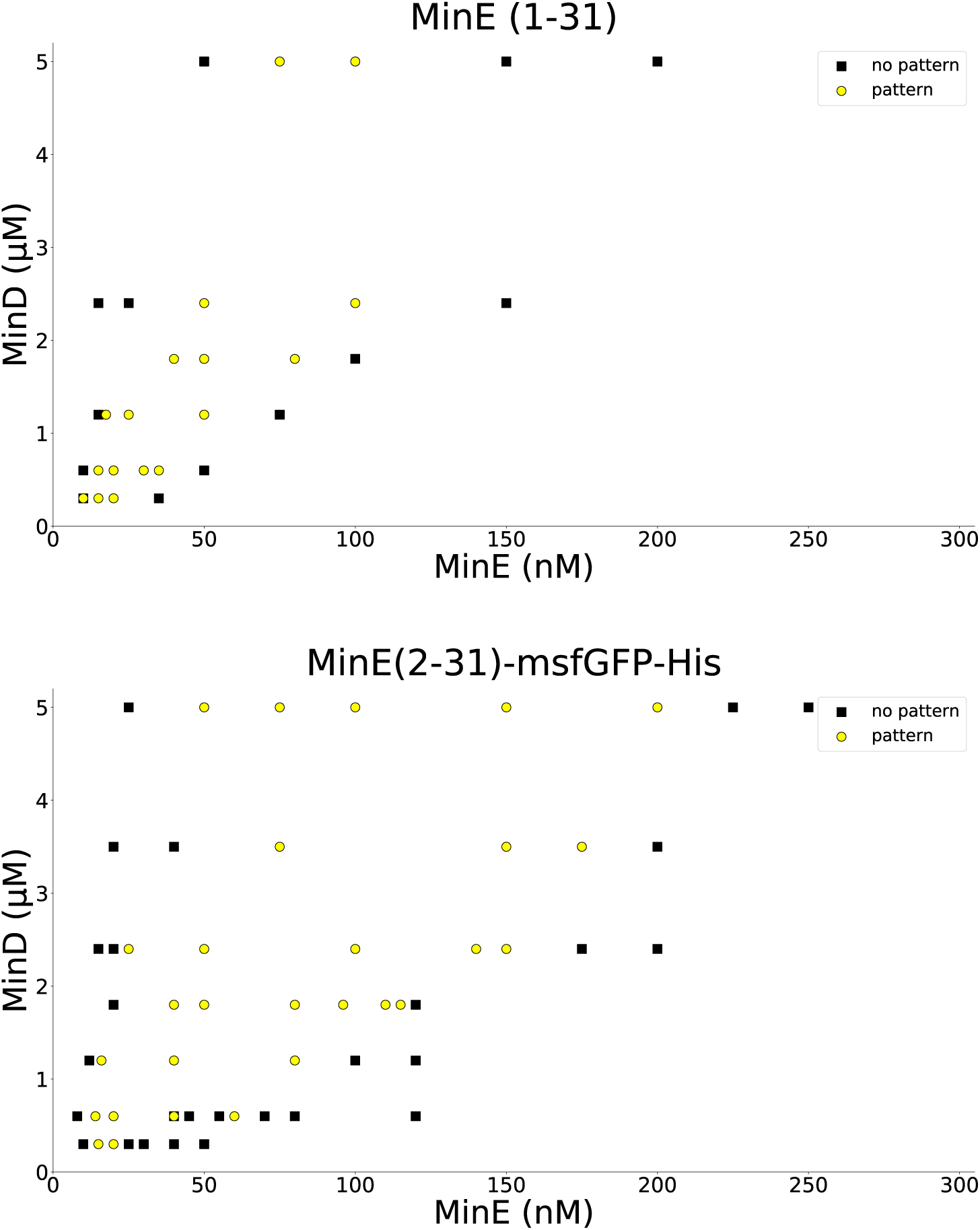
MinD and the peptide MinE(1-31) or MinE(2-31)-sfGFP, respectively, were titrated to find the range in which patterns are formed. All experiments were done on SLBs consisting of DOPC:DOPG (2:1). Similar titrations for full-length MinE can be found in (***Glock et al., 2018b***). Wild-type MinE generally forms patterns with MinD in a much larger range, going beyond 10 µM.

**Figure 3–Figure supplement 1.**
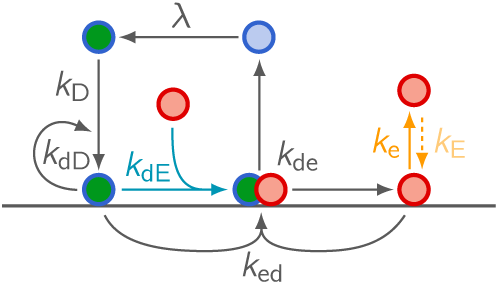
Network cartoon of the MinE “skeleton” model extended by MinE membrane binding.

**Figure 3–Figure supplement 2.**
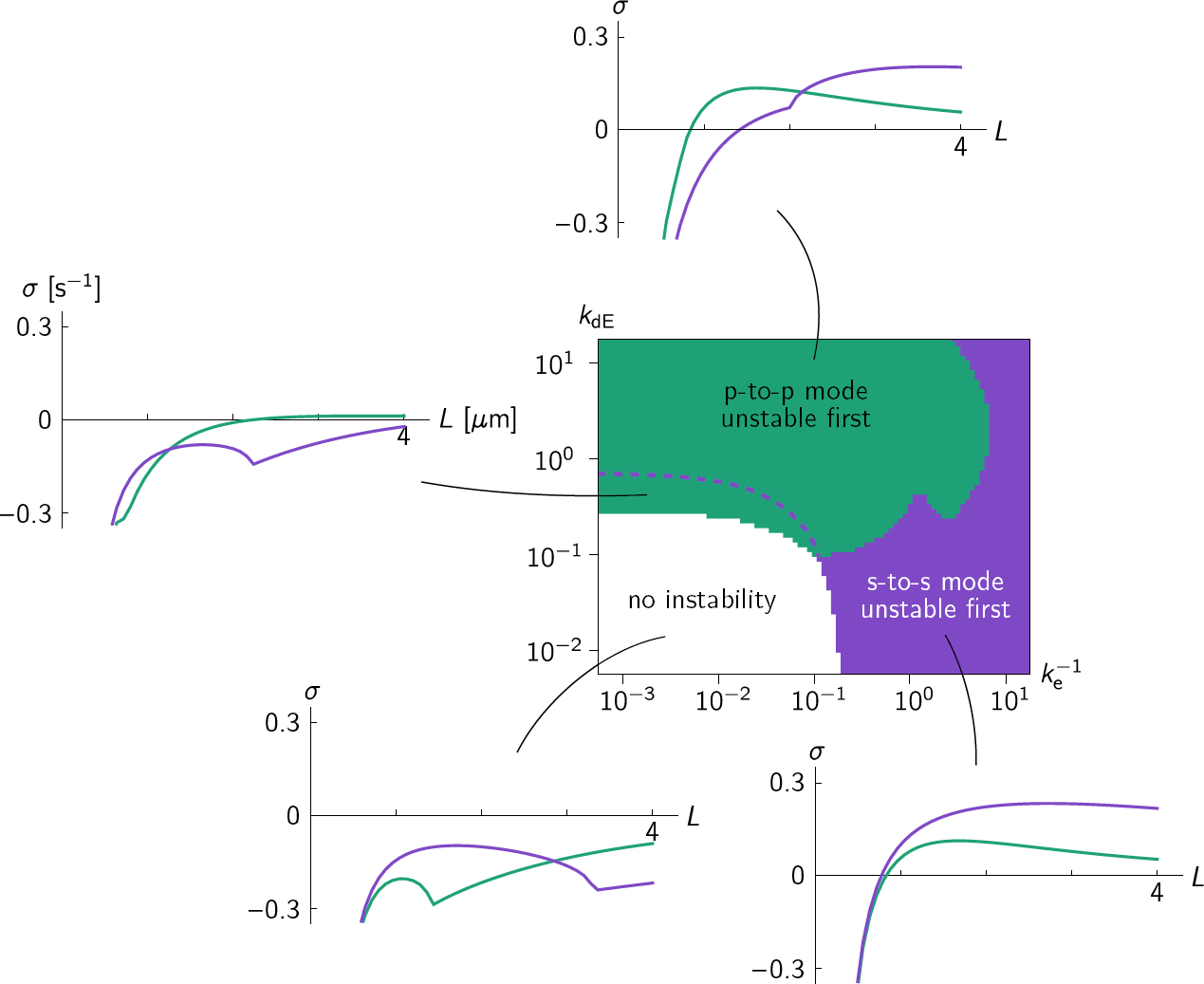
Linear stability analysis in the ellipse geometry. Regions in the phase diagram are colored according to which eigenmode (green for pole-to-pole, purple for side-to-side mode) becomes unstable first for increasing cell size. Above the dashed purple line, the side-to-side mode grows faster at grown cell size (*L* = 4 µm). Typical relationships between cell size *L* and growth rate *σ* of the pole-to-pole mode (green line) and side-to-side mode (purple line) are shown for each parameter region. Comparison to the phase diagram from numerical simulations (**Figure 3**(b)) shows that the mode becoming unstable first, not the fastest growing mode at full cell size, predicts the axis selected by the fully developed pattern.

## References

Caspi Y, Dekker C. Mapping out Min protein patterns in fully confined fluidic chambers. eLife. 2016 nov; 5(iii):1–53. http://elifesciences.org/lookup/doi/10.7554/eLife.19271, doi: 10.7554/eLife.19271.

Denk J, Kretschmer S, Halatek J, Hartl C, Schwille P, Frey E. MinE conformational switching confers robustness on self-organized Min protein patterns. Proceedings of the National Academy of Sciences of the United States of America. 2018 may; 115(18):4553–4558.http://www.pnas.org/lookup/doi/10.1073/pnas.1719801115http://www.ncbi.nlm.nih.gov/pubmed/29666276http://www.pubmedcentral.nih.gov/articlerender.fcgi?artid=PMC5939084,doi: 10.1073/pnas.1719801115.

Downing BPB, Rutenberg AD, Touhami A, Jericho M. Subcellular Min oscillations as a single-cell reporter of the action of polycations, protamine, and gentamicin on Escherichia coli. PloS one. 2009 sep; 4(9):e7285. http://www.ncbi.nlm.nih.gov/pubmed/19789705http://www.pubmedcentral.nih.gov/articlerender.fcgi?artid=PMC2749335, doi: 10.1371/journal.pone.0007285.

Ernst O, Zor T. Linearization of the Bradford Protein Assay. Journal of Visualized Experiments. 2010 apr; (38):1–6. http://www.jove.com/index/Details.stp?ID=1918 http://www.ncbi.nlm.nih.gov/pubmed/20386536http://www.pubmedcentral.nih.gov/articlerender.fcgi?artid=PMC3164080, doi: 10.3791/1918.

Gessele R, Halatek J, Frey E. PAR protein activation-deactivation cycles stabilize long-axis polarization in C. elegans. bioRxiv. 2018; https://www.biorxiv.org/content/early/2018/10/25/451880, doi: 10.1101/451880.

Glock P, Broichhagen J, Kretschmer S, Blumhardt P, Mücksch J, Trauner D, Schwille P. Optical Control of a Biological Reaction-Diffusion System. Angewandte Chemie International Edition. 2018 feb; 57(9):2362–2366. http://doi.wiley.com/10.1002/anie.201712002http://www.ncbi.nlm.nih.gov/pubmed/29266672, doi: 10.1002/anie.201712002.

Glock P, Ramm B, Heermann T, Kretschmer S, Schweizer J, Mücksch J, Alagöz G, Schwille P. Stationary patterns in a two-protein reaction-diffusion system. ACS Synthetic Biology. 2018; p. acssynbio.8b00415. http://pubs.acs.org/doi/10.1021/acssynbio.8b00415, doi: 10.1021/acssynbio.8b00415.

Green JBA, Sharpe J. Positional information and reaction-diffusion: two big ideas in developmental biology combine. Development. 2015 apr; 142(7):1203–1211. http://dev.biologists.org/cgi/doi/10.1242/dev.114991http://www.ncbi.nlm.nih.gov/pubmed/25804733, doi: 10.1242/dev.114991.

Halatek J, Frey E. Rethinking pattern formation in reaction–diffusion systems. Nature Physics. 2018 may; 14(5):507–514. http://dx.doi.org/10.1038/s41567-017-0040-5http://www.nature.com/articles/349s41567-017-0040-5,doi: 10.1038/s41567-017-0040-5.

Halatek J, Brauns F, Frey E. Self-organization principles of intracellular pattern formation. Philosophical transactions of the Royal Society of London Series B, Biological sciences. 2018 may; 373(1747). http://dx.doi.org/10.1098/rstb.2017-0107http://www.ncbi.nlm.nih.gov/pubmed/29632261http://arxiv.org/abs/1802.07169{%}0Aw http://dx.doi.org/10.1098/rstb.2017-0107, doi: 10.1098/rstb.2017.0107.

Halatek J, Frey E. Highly Canalized MinD Transfer and MinE Sequestration Explain the Origin of Robust MinCDE-Protein Dynamics. Cell Reports. 2012; 1(6):741–752. http://dx.doi.org/10.1016/j.celrep.2012.04.005, doi: 10.1016/j.celrep.2012.04.005.

Hu Z, Gogol EP, Lutkenhaus J. Dynamic assembly of MinD on phospholipid vesicles regulated by ATP and MinE. Proceedings of the National Academy of Sciences of the United States of America. 2002 may; 99(10):6761–6. http://www.ncbi.nlm.nih.gov/pubmed/11983867 http://www.pubmedcentral.nih.gov/articlerender.fcgi?artid=PMC124476, doi: 10.1073/pnas.102059099.

Hu Z, Lutkenhaus J. Topological regulation of cell division in E. coli: Spatiotemporal oscillation of MinD requires stimulation of its ATPase by MinE and phospholipid. Molecular Cell. 2001; 7(6):1337–1343. doi: 10.1016/S1097-2765(01)00273-8.

Huang KC, Meir Y, Wingreen NS. Dynamic structures in Escherichia coli: spontaneous formation of MinE rings and MinD polar zones. Proceedings of the National Academy of Sciences of the United States of America. 2003 oct; 100(22):12724–8. http://www.ncbi.nlm.nih.gov/pubmed/14569005 http://www.pubmedcentral.nih.gov/articlerender.fcgi?artid=PMC240685, doi: 10.1073/pnas.2135445100.

Ivanov V, Mizuuchi K. Multiple modes of interconverting dynamic pattern formation by bacterial cell division proteins. Proceedings of the National Academy of Sciences of the United States of America. 2010; 107(18):8071–8078. doi: 10.1073/pnas.0911036107.

Kretschmer S, Zieske K, Schwille P. Large-scale modulation of reconstituted Min protein patterns and gradients by de1ned mutations in MinE’s membrane targeting sequence. PLoS ONE. 2017; p. 1–16. doi:https://doi.org/10.1371/journal.pone.0179582.

Lackner LL, Raskin DM, De Boer PaJ. ATP-dependent interactions between Escherichia coli Min proteins and the phospholipid membrane in vitro. Journal of Bacteriology. 2003; 185(3):735–749. doi: 10.1128/JB.185.3.735-749.2003.

Loose M, Fischer-Friedrich E, Herold C, Kruse K, Schwille P. Min protein patterns emerge from rapid rebinding and membrane interaction of MinE. Nature structural & molecular biology. 2011; 18(5):577–583. http://dx.doi.org/10.1038/nsmb.2037, doi: 10.1038/nsmb.2037.

Loose M, Fischer-Friedrich E, Ries J, Kruse K, Schwille P. Spatial regulators for bacterial cell division self-organize into surface waves in vitro. Science (New York, NY). 2008; 320(5877):789–792. doi: 10.1126/science.1154413.

O’Shea E, Rutkowski R, Stafford W, Kim P. Preferential heterodimer formation by isolated leucine zippers from fos and jun. Science. 1989 aug; 245(4918):646–648. http://www.sciencemag.org/cgi/doi/10.1126/science.2503872http://www.ncbi.nlm.nih.gov/pubmed/2503872, doi: 10.1126/science.2503872.

Park KT, Wu W, Battaile KP, Lovell S, Holyoak T, Lutkenhaus J. The min oscillator uses MinD-dependent conformational changes in MinE to spatially regulate cytokinesis. Cell. 2011; 146(3):396–407. http://dx.doi.org/10.1016/j.cell.2011.06.042, doi: 10.1016/j.cell.2011.06.042.

Pichoff S, Vollrath B, Touriol C, Bouché JP. Deletion analysis of gene minE which encodes the topological speci1city factor of cell division in Escherichia coli. Molecular microbiology. 1995; 18(2):321–329.

Ramm B, Glock P, Schwille P. In Vitro Reconstitution of Self-Organizing Protein Patterns on Supported Lipid Bilayers. Journal of Visualized Experiments. 2018 jul; (137):e58139. https://www.jove.com/video/58139https://www.jove.com/video/58139/in-vitro-reconstitution-self-organizing-protein-patterns-on-supported, doi: 10.3791/58139.

Szalóki N, Krieger JW, Komáromi I, Tóth K, Vámosi G. Evidence for Homodimerization of the c-Fos Transcription Factor in Live Cells Revealed by Fluorescence Microscopy and Computer Modeling. Mol Cell Biol. 2015; 35(August):3785–3798. http://mcb.asm.org/content/35/21/3785.abstract{%}5Cnalsdb.org, doi: 10.1128/MCB.00346-15.

Szeto TH, Rowland SL, Habrukowich CL, King GF. The MinD membrane targeting sequence is a transplantable lipid-binding helix. Journal of Biological Chemistry. 2003; 278(41):40050–40056. doi: 10.1074/jbc.M306876200.

Vecchiarelli AG, Li M, Mizuuchi M, Hwang LC, Seol Y, Neuman KC, Mizuuchi K. Membrane-bound MinDE complex acts as a toggle switch that drives Min oscillation coupled to cytoplasmic depletion of MinD. Proceedings of the National Academy of Sciences. 2016 mar; 113(11):E1479–E1488. http://www.pnas.org/lookup/doi/10.1073/pnas.1600644113, doi: 10.1073/pnas.1600644113.

Vecchiarelli AG, Li M, Mizuuchi M, Mizuuchi K. Differential aZnities of MinD and MinE to anionic phospholipid influence Min patterning dynamics in vitro. Molecular Microbiology. 2014; 93(3):453–463. doi: 10.1111/mmi.12669.

Wu F, Halatek J, Reiter M, Kingma E, Frey E, Dekker C. Multistability and dynamic transitions of intracellular Min protein patterns. Molecular Systems Biology. 2016; 12(6):873. http://msb.embopress.org/lookup/doi/10.5252/msb.20156724, doi: 10.15252/msb.20156724.

